# UFMylation of 14-3-3ε coordinates MAVS signaling complex assembly to promote antiviral innate immune induction

**DOI:** 10.1101/2025.03.19.644084

**Authors:** Isaura Vanessa Gutierrez, Moonhee Park, Lauren Sar, Ryan E. Rodriguez, Hannah M. Schmidt, Daltry L. Snider, Gabriella Torres, K. Matthew Scaglione, Stacy M. Horner

**Affiliations:** Department of Integrative Immunobiology, Duke University School of Medicine, Durham, NC, USA; Department of Molecular Genetics and Microbiology, Duke University School of Medicine, Durham, NC, USA; Duke Center for Neurodegeneration and Neurotherapeutics, Duke University School of Medicine, Durham, NC, USA; Department of Medicine, Duke University School of Medicine, Durham, NC, USA

**Author notes:** Correspondence Stacy M. Horner, Ph.D, Duke University School of Medicine 213 Research Dr., Box 3053 DUMC Durham, NC USA 27710.

## Abstract

Post-translational modifications regulate RIG-I signaling in diverse ways. We previously showed that UFMylation, the covalent attachment of the ubiquitin-fold modifier UFM1 to proteins, enhances RIG-I signaling by promoting its interaction with its membrane-targeting adaptor 14-3-3ε. Here, we map UFM1 conjugation to lysines K50 and K215 on 14-3-3ε and demonstrate how these UFMylation events control RIG-I signaling. Using *in vitro* and cellular UFMylation assays, we reveal that K50R/K215R mutations abolish UFMylation and reduce type I and III interferon induction following RIG-I activation. Unexpectedly, these mutations do not disrupt 14-3-3ε-RIG-I interaction. Instead, they paradoxically enhance RIG-I interaction with MAVS while simultaneously reducing 14-3-3ε-MAVS interaction. These findings establish UFMylation of 14-3-3ε as an important control that shapes MAVS complex architecture to ensure optimal RIG-I signaling and highlights the broader regulatory role of UFMylation in antiviral innate immunity.

**Importance:** Post-translational modifications provide regulatory control of antiviral innate immune responses. Our study reveals that UFMylation of 14-3-3ε is required for RIG-I-mediated innate immune signaling. We demonstrate that conjugation of UFM1 to specific lysine residues on 14-3-3ε enhances downstream signaling events that facilitate interferon induction. It does this by stabilizing 14-3-3ε association with the MAVS signaling complex and coordinating productive complex architecture. By identifying the precise sites of UFMylation on 14-3-3ε and their functional consequences, we provide insights into the regulatory layers governing antiviral innate immunity. These findings complement emerging evidence that UFMylation serves as a versatile modulator across diverse immune pathways. Furthermore, our work highlights how protein chaperones like 14-3-3ε can be dynamically modified to orchestrate complex signaling cascades, suggesting potential therapeutic approaches for targeting dysregulated innate immunity.

## Introduction

The antiviral innate immune response is tightly regulated to ensure effective defense against viral infections while preventing excessive inflammation. This antiviral response to RNA viruses is activated by cytosolic RIG-I-like receptors (RLRs) RIG-I and MDA5, which detect pathogen-associated molecular patterns from RNA viruses and initiate the formation of a MAVS-signaling complex (1). Upon activation, MAVS undergoes oligomerization and recruits downstream signaling proteins to form a large multiprotein assembly known as the MAVS signalosome, which serves as a platform for signal amplification and propagation. This signaling cascade ultimately leads to the transcriptional activation of type I and type III interferons (IFN) and the production of IFN-stimulated genes (ISGs) that limit viral infection. Robust innate immune signaling is essential for effective adaptive immunity, but dysregulation of these pathways can result in autoimmune diseases and interferonopathies (2–4). Therefore, the induction of IFN is carefully regulated by mechanisms that temporarily activate, dampen, and ultimately shut down the signaling pathway.

Both 14-3-3ε and 14-3-3η are RLR regulatory proteins that control IFN induction. These proteins facilitate the translocation of RIG-I and MDA5, respectively, from the cytosol to intracellular membrane signaling sites to interact with the MAVS signalosome (5, 6). 14-3-3 proteins are highly conserved scaffolding proteins involved in diverse cellular processes, including signal transduction, apoptosis, and stress responses. Typically, they coordinate diverse cellular processes by binding to phosphorylated client proteins, although they are also known to bind to non-phosphorylated client proteins (7–13). Recent work has also suggested that 14-3-3 proteins can serve as protein chaperones, preventing aggregation and phase separation of their client proteins (14). The essential role of 14-3-3ε and 14-3-3η in antiviral innate immunity is underscored by the observation that multiple viruses have evolved specific mechanisms to antagonize these proteins. For example, orthoflavivirus NS3 proteins specifically target and inactivate both 14-3-3ε and 14-3-3η to prevent RLR signaling, the enterovirus 3C protease cleaves 14-3-3ε, and the influenza A virus NS1 protein binds to 14-3-3ε (15–18). These diverse viral targeting strategies highlight how critical 14-3-3 proteins are for mounting an antiviral innate immune response.

Post-translational modifications play a crucial role in regulating RIG-I signaling (19, 20). Recent studies by our group have shown that ubiquitin-fold modifier 1 (UFM1) and the proteins involved in UFM1 conjugation promote RIG-I signaling (21). UFMylation involves the covalent conjugation of ubiquitin-fold modifier 1 (UFM1), an 85-amino acid peptide, to lysine residues on proteins, utilizing an E1 activating enzyme (UBA5), an E2 conjugating enzyme (UFC1), and an E3 scaffold-like ligase complex (UFL1 and the membrane-anchored protein UFBP1) (22–30). UFM1 is a non-degradative mark, and its addition to proteins can promote protein stability by antagonizing ubiquitination or stabilizing interactions with other proteins (31–33). UFMylation modulates various cellular processes, including the DNA damage response and genomic instability, various aspects of ER homeostasis and translation, ER-Golgi transport, viral infection, cancer progression, and both adaptive and innate immune responses (24, 25, 27, 28, 33–46).

In the context of innate immunity, UFMylation plays diverse regulatory roles. For example, UFMylation promotes NLRP3 inflammasome activation (47), while in other contexts it can limit LPS-mediated inflammation via its roles in regulating ER homeostasis and the unfolded protein response (48–51). Additionally, UFL1 can bind STING during herpes simplex virus 1 infection to maintain IFN induction (52). However, the overall relationship between UFMylation and viral infection is complex. In some cases, UFMylation can promote viral infection. For example, UFMylation of the ribosomal protein RPL26 enhances hepatitis A virus translation (53), and in orthoflaviviruses, UFMylation promotes the production of infectious virions (54). Reflecting the importance of UFMylation in host defense, viruses have evolved mechanisms to manipulate host UFMylation pathways by targeting UFL1 to specific substrates. For example, orthoflaviviruses co-opt UFL1 to aid in viral assembly (54), while Epstein-Barr virus targets UFL1 to MAVS for UFMylation to dampen inflammasome activation (55). These diverse roles highlight UFMylation as a key regulatory system in host-pathogen interactions.

Our previous work on UFMylation in the context of RIG-I signaling revealed that UFL1 translocates to intracellular membranes upon RIG-I activation and that UFL1 and UFM1 broadly facilitate RIG-I interaction with 14-3-3ε, with 14-3-3ε exhibiting increased UFMylation in response to RIG-I activation (21, 56). In this study, we established the molecular mechanism by which UFMylation of 14-3-3ε promotes RIG-I signaling. Using cellular and *in vitro* UFMylation assays, we identify lysine residues K50 and K215 as the mono-UFMylation sites of 14-3-3ε. Through mutagenesis studies, we demonstrate that preventing UFMylation at these residues by introducing K50R/K215R mutations impairs both type I and type III interferon induction in response to RIG-I activation. Unexpectedly, these UFMylation-deficient mutations do not disrupt the 14-3-3ε-RIG-I interaction. Instead, UFMylation of 14-3-3ε promotes signaling downstream of RIG-I recruitment by stabilizing an interaction between 14-3-3ε and MAVS. In the absence of 14-3-3ε UFMylation, the 14-3-3ε-MAVS interaction is weakened while RIG-I-MAVS binding is paradoxically enhanced, suggesting that UFMylated 14-3-3ε coordinates optimal MAVS signalosome assembly to enable robust interferon production. These findings reveal that UFMylation of 14-3-3ε itself promotes signaling from the RIG-I-MAVS complex, highlighting the broader role of UFMylation in antiviral innate immunity.

## Results

### UFM1 conjugation is required to promote 14-3-3ε-mediated enhancement of RIG-I signaling to IFN-β

To define the molecular mechanism by which UFMylation to 14-3-3ε specifically regulates RIG-I signaling, we first established the functional requirement for UFM1 and its conjugation activity in signaling to the IFN-β promoter. To test if UFM1 enhances RIG-I signaling through 14-3-3ε, we performed IFN-β promoter luciferase assays in 293T cells following ectopic expression of UFM1, 14-3-3ε, and RIG-I. We found that UFM1 ectopic expression significantly enhanced RIG-I driven signaling to the IFN-β promoter. This signaling was further amplified when UFM1 was co-expressed with both RIG-I and 14-3-3ε (Figure 1A). Next, to determine whether UFM1-conjugation is required for this enhancement of RIG-I signaling, we generated a conjugation-deficient UFM1 mutant by deleting three amino acids from its C-terminus to prevent covalent attachment of UFM1 to target proteins (ΔC3) (22). Importantly, both WT and conjugation-deficient UFM1 are expressed at similar levels in cells (Figure 1B). Using IFN-β promoter luciferase assays in 293T cells, we found that unlike wild-type UFM1, the conjugation-deficient UFM1 mutant failed to enhance 14-3-3ε-driven RIG-I signaling (Figure 1B). These results demonstrate that UFM1 must be conjugation-competent to facilitate 14-3-3ε-mediated enhancement of RIG-I signaling to the IFN-β promoter.

**Figure 1.**
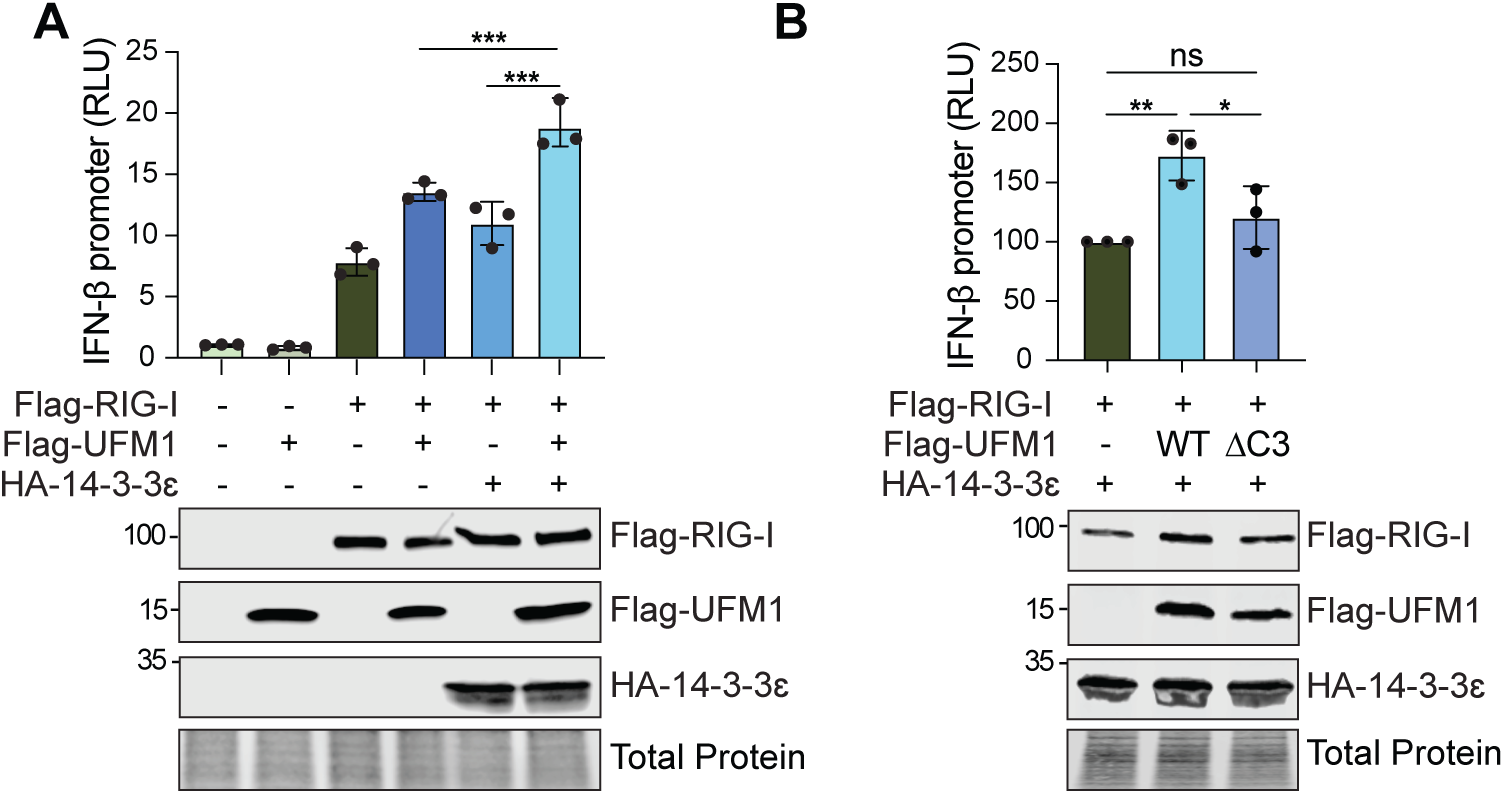
UFM1 conjugation is required to promote 14-3-3ε-mediated enhancement of RIG-I signaling to IFN-β. (A) IFN-β promoter reporter luciferase expression (relative to *Renilla)* from 293T cells expressing Vector, Flag-UFM1^WT^, Flag-RIG-I, or HA-14-3-3ε. (B) IFN-β promoter reporter luciferase expression (relative to *Renilla*) from 293T cells expressing Vector, Flag-UFM1^WT^, Flag-UFM1^ΔC3^, Flag-RIG-I, or HA-14-3-3ε. Bars represent the mean ± SD, *n =* 3 independent replicates. *p ≤ 0.05, ** ≤ 0.01, ***p ≤ 0.001 determined by one way ANOVA followed by Šidák’s multiple comparisons test (A) or Tukey’s multiple comparisons test (B).

### UFM1 conjugation to 14-3-3ε requires residues K50 and K215

Having established that active UFM1 conjugation is required for 14-3-3ε to enhance RIG-I signaling, we next sought to verify our previous published results that 14-3-3ε is directly UFMylated (21) and determine which specific lysine residues serve as UFM1-conjugation sites. First, we verified 14-3-3ε UFMylation in cells. For this, we developed a modified UFMylation assay based on previous studies (57). Because UFMylation can be low stoichiometry and transient, overexpression of pathway components and the use of a deconjugation-resistant UFM1 (G83A) are typically used to reveal substrate modification. Thus, we co-expressed HA-tagged 14-3-3ε, Flag-UFM1 (WT, G83A, or ΔC3), and the UFMylation ligase complex proteins Myc-UFL1 and Myc-UFBP1 in 293T-UFM1 KO cells (21). Critically, lysates were boiled prior to immunoprecipitation to denature protein complexes, ensuring that only proteins covalently linked to the immunoprecipitated protein would be detected in subsequent analyses. Following anti-HA immunoprecipitation under denaturing conditions, we identified a single apparent band with a shift of approximately 15 kDa, consistent with predominantly a mono-UFMylation event on 14-3-3ε (Figure 2A). Importantly, the conjugation-deficient Flag-UFM1-ΔC3 showed no attachment of 14-3-3ε, while a deconjugation-resistant mutant, Flag-UFM1-G83A (22), demonstrated significantly increased levels of 14-3-3ε UFMylation.

**Figure 2.**
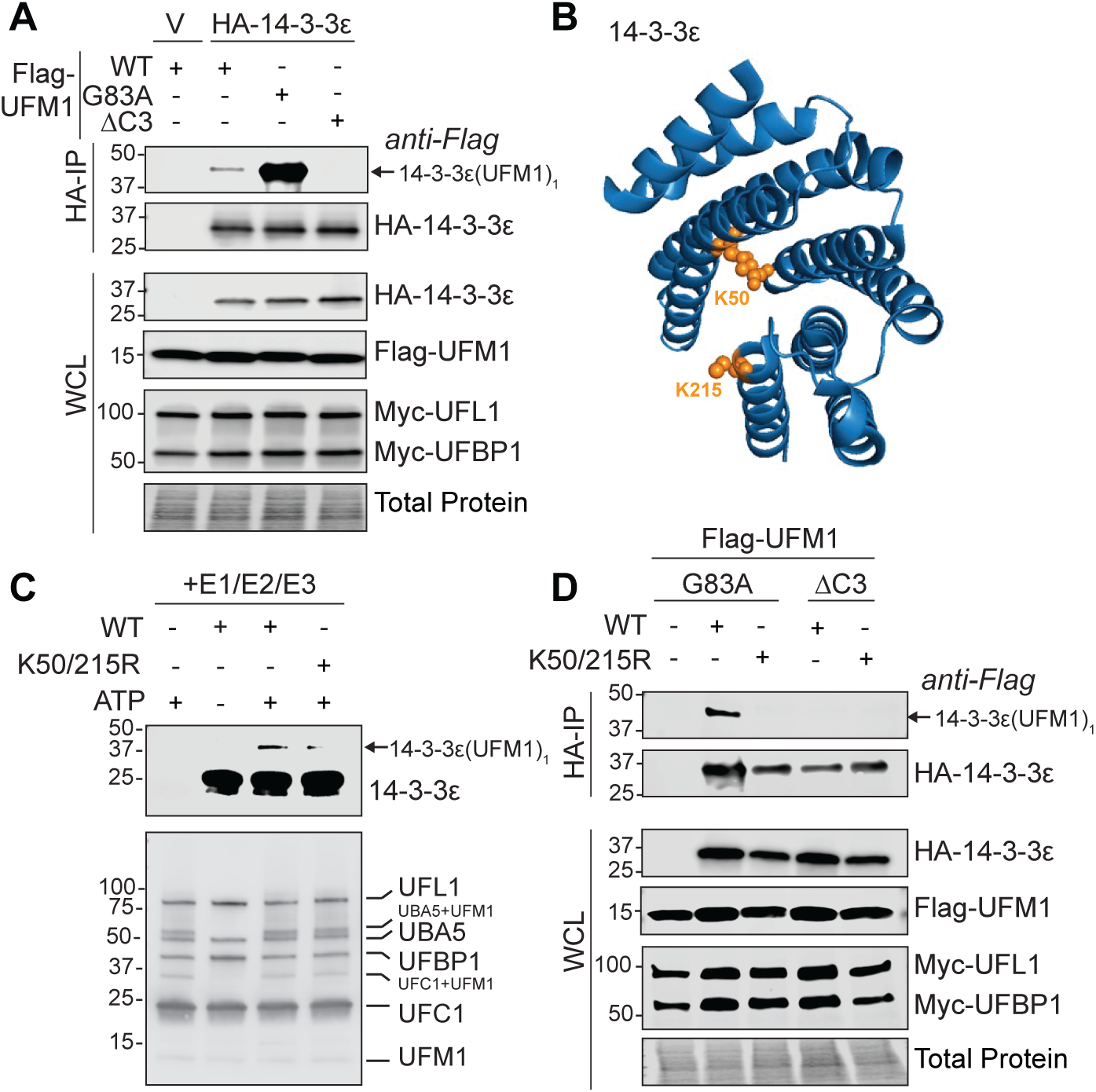
UFM1 conjugation to 14-3-3ε requires residues K50 and K215 on 14-3-3ε. (A) UFMylation assay of anti-HA immunoprecipitated lysates from 293T UFM1 KO cells expressing Vector (V) or HA-14-3-3ε along with Myc-UFL1, Myc-UFBP1, and Flag-UFM1-WT, -G83A, or -ΔC3. (B) Ribbon representation of 14-3-3ε (PDB entry 3UAL), with amino acids K50 and K215 indicated in orange. (C) Purified UFMylation proteins (UFBP1, UFL1, UFC1, UBA5, and UFM1) as well as 14-3-3ε^WT^ or 14-3-3ε^K50/215R^ were incubated in UFMylation assay buffer in the presence or absence of ATP, followed by immunoblot analysis for 14-3-3ε (top) or using an antibody cocktail against all UFMylation proteins (bottom). (D) UFMylation assay of anti-HA immunoprecipitated lysates from 293T UFM1 KO cells expressing Vector (-), Flag-UFM1-WT, -G83A, or -ΔC3, and HA-14-3-3ε^WT^ or HA-14-3-3ε^K50/215R^. Representative blots from *n =* 3 independent replicates are shown (A, C, and D).

We next sought to identify the specific lysine residues on 14-3-3ε that undergo UFMylation by using mass spectrometry analysis. We used Huh7 cells stably expressing Flag-UFM1 with a two amino acid deletion (ΔC2). This mutant enhances detection of UFMylated proteins by eliminating the prerequisite UFSP1/UFSP2-mediated cleavage of the last two amino acids in UFM1 required to initiate UFM1 conjugation, (22). After immunoprecipitation of HA-14-3-3ε, mass spectrometry analysis suggested that two lysine residues, K50 and K215, were covalently modified by UFM1. Both K50 and K215 reside within clusters of basic residues within a positively charged pocket of the conserved amphipathic ligand binding groove of 14-3-3 proteins (Figure 2B) (58). This groove mediates interactions with phosphorylated and non-phosphorylated client proteins, suggesting that UFMylation at these sites could directly modulate 14-3-3ε protein-protein interactions.

To confirm these specific sites as bona fide UFMylation targets, we employed both *in vitro* and cellular UFMylation assays. Using an *in vitro* UFMylation assay with purified UFMylation proteins (UBA5, UFC1, UFBP1/UFL1, and UFM1), we found that wild-type 14-3-3ε is mono-UFMylated, whereas lysine-to-arginine (K to R) mutations at K50 and K215 (K50R/K215R) abolished this mono-UFMylation (Figure 2C). Similarly, in the cellular UFMylation assay, the K50R-K215R mutations significantly abrogated 14-3-3ε UFMylation (Figure 2D). These complementary approaches confirm that K50 and K215 are necessary for detectable mono-UFMylation of 14-3-3ε both *in vitro* and in cell-based assays. However, as we only detect mono-UFMylation of 14-3-3ε, these results indicate that each individual 14-3-3ε molecule is UFMylated at either K50 or K215, but not both simultaneously. This is consistent with the structural positioning of these residues on opposite faces of the ligand-binding groove (58). We note that the relative occupancy and functional contribution of each site remain to be determined and may vary under different cellular conditions. Therefore, we used the K50R/K215R double point mutant for further exploration of the biological impacts of 14-3-3ε UFMylation.

### K50 and K215 are required for 14-3-3ε-mediated RIG-I activation of IFN

Having identified K50 and K215 as the UFMylation sites on 14-3-3ε, we next investigated whether these residues are functionally required for 14-3-3ε to enhance RIG-I-mediated signaling. We used two complementary approaches to assess the impact of UFMylation-deficient 14-3-3ε on interferon induction. First, we evaluated IFN-β promoter activation using a luciferase reporter assay in 293T-RIG-I KO cells (59). We used these cells to establish a reconstitution system where RIG-I signaling is specifically derived from co-expressed constructs, eliminating confounding effects from endogenous RIG-I. Importantly, RIG-I ectopic expression can be sufficient to drive signaling. We ectopically expressed 14-3-3ε^WT^ or 14-3-3ε^K50/215R^, along with RIG-I, and measured luciferase activity (Figure 3A). While 14-3-3ε^WT^ significantly increased RIG-I activation of the IFN-β promoter, as expected, 14-3-3ε^K50/215R^ displayed a significantly impaired ability to enhance this signaling (Figure 3A). This result suggests that UFMylation at K50 or K215 is critical for 14-3-3ε to promote RIG-I-mediated transcriptional activation of the IFN-β promoter.

**Figure 3.**
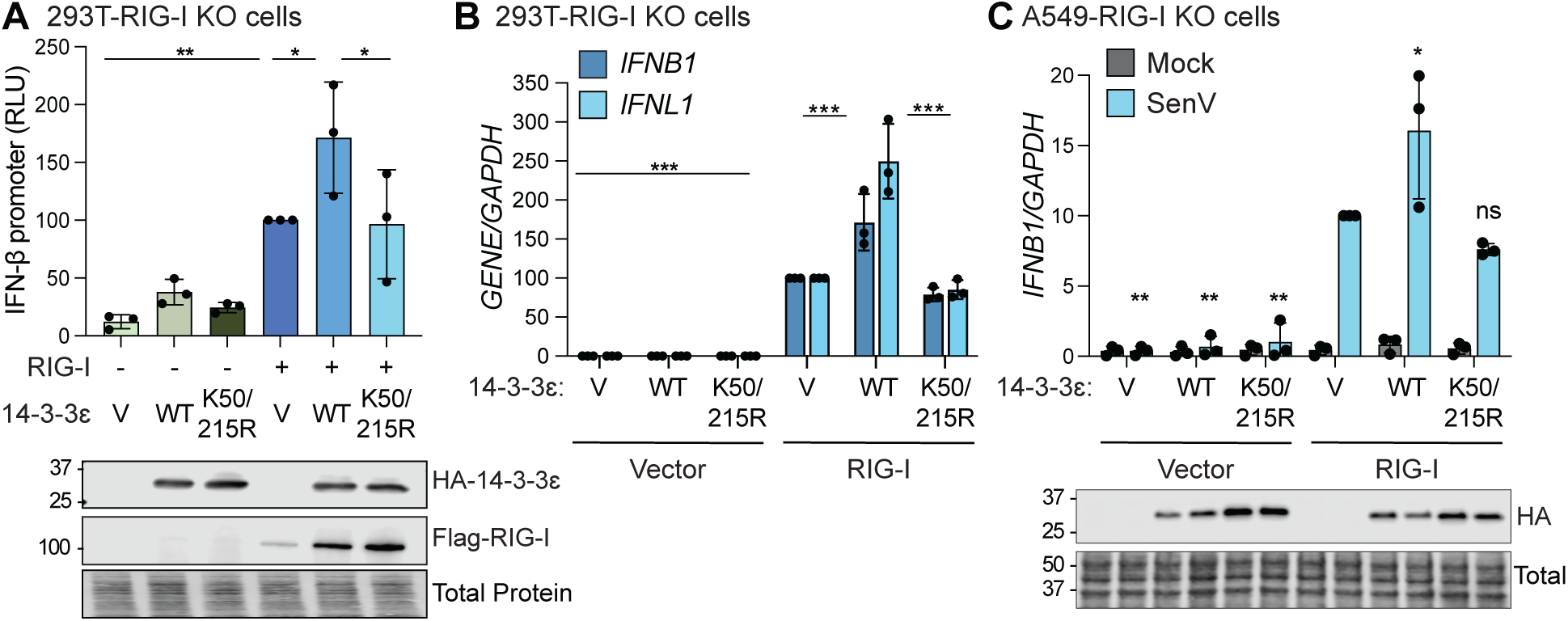
K50 and K215 are required for 14-3-3ε-mediated RIG-I activation of IFN. (A) IFN-β promoter reporter *Gaussia* luciferase expression from 293T-RIG-I KO cells expressing Vector (V), Flag-RIG-I, and HA-14-3-3ε^WT^ or HA-14-3-3ε^K50/215R^, with samples expressing RIG-I + V normalized to 100. (B) RT-qPCR analysis (rel. to *GAPDH*) of RNA extracted from 293T-RIG-I KO cells transfected with Flag-RIG-I, and HA-14-3-3ε^WT^ or HA-14-3-3ε^K50/215R^ followed by SenV infection (6 h), with RIG-I + V samples normalized to 100. (C) RT-qPCR analysis (rel. to *GAPDH*) of RNA extracted from A549-RIG-I KO cells transfected with low amounts of Flag-RIG-I, and HA-14-3-3ε^WT^ or HA-14-3-3ε^K50/215R^, followed by SenV infection (6 h), with RIG-I +V samples normalized to 100. *p ≤ 0.05, ** ≤ 0.01, ***p ≤ 0.005 determined by one way ANOVA followed by (A) Bonferroni or (B-C) Tukey’s multiple comparisons test. Mean ± SD, *n =* 3 independent replicates are shown.

To validate these findings using a complementary approach, we measured endogenous interferon transcript levels by RT-qPCR. We co-expressed RIG-I with 14-3-3ε^WT^ or 14-3-3ε^K50/215R^ in 293T-RIG-I KO cells and infected them with Sendai virus (SenV). SenV is an RNA virus that is a potent activator of RIG-I signaling (60, 61). Importantly, SenV is sensed specifically by RIG-I, ensuring our findings reflect RIG-I-driven signaling. Neither 14-3-3ε induced basal *IFNB1* and *IFNL1* expression without RIG-I. However, in the presence of RIG-I, 14-3-3ε^WT^ enhanced interferon transcript levels while 14-3-3ε^K50/215R^ did not (Figure 3B), confirming the luciferase results. To validate these findings in another cell type, we measured the RNA levels of *IFNB1* in A549-RIG-I KO cells, generated by CRISPR/Cas9, in response to SenV infection. Here, we co-expressed RIG-I with 14-3-3ε^WT^ or 14-3-3ε^K50/215R^. Consistent with our earlier results, we found that 14-3-3ε^WT^ enhances RIG-I-mediated induction of *IFNB1* in response to SenV, while 14-3-3ε^K50/215R^ does not (Figure 3C). Collectively, these findings demonstrate K50 and K215 are critical for 14-3-3ε to promote transcriptional activation of type I and type III IFNs in response to RIG-I activation, suggesting that UFMylation enhances RIG-I-mediated induction of IFN.

### K50 and K215 are not required for 14-3-3ε interaction with RIG-I

Having established that K50 and K215 are required for 14-3-3ε to promote RIG-I signaling, we next investigated the mechanism underlying this functional requirement. Since our previous findings revealed that both UFM1 and UFL1 promote the interaction of RIG-I with 14-3-3ε (21), we hypothesized that UFMylation of 14-3-3ε at K50/K215 would be required for its interaction with RIG-I during SenV-mediated RIG-I activation. To test this hypothesis, we performed co-immunoprecipitation experiments comparing the interaction between RIG-I and either 14-3-3ε^WT^ or the UFMylation-deficient 14-3-3ε^K50/215R^ mutant. Unexpectedly, we found that 14-3-3ε^K50/215R^ maintained interaction with RIG-I at levels comparable to the wild-type protein (Figure 4A-B), indicating that UFMylation at these residues is not essential for the association of 14-3-3ε and RIG-I. To validate our experimental approach, we included a control mutant of 14-3-3ε where key residues at the C-terminal end (EDE to AAA) were altered, as these previously have been reported to mediate this interaction (62). As expected, this control mutant showed significantly reduced RIG-I association, confirming the work of others, as well as the specificity of the assay. These results revealed that UFMylation at K50 or K215 are not critical for the interaction between 14-3-3ε and RIG-I. Further, since 14-3-3ε^K50/215R^ retains RIG-I binding comparable to wild-type, the signaling defects observed in Figure 3 are unlikely to result from global disruptions to the 14-3-3ε ligand-binding groove.

**Figure 4.**
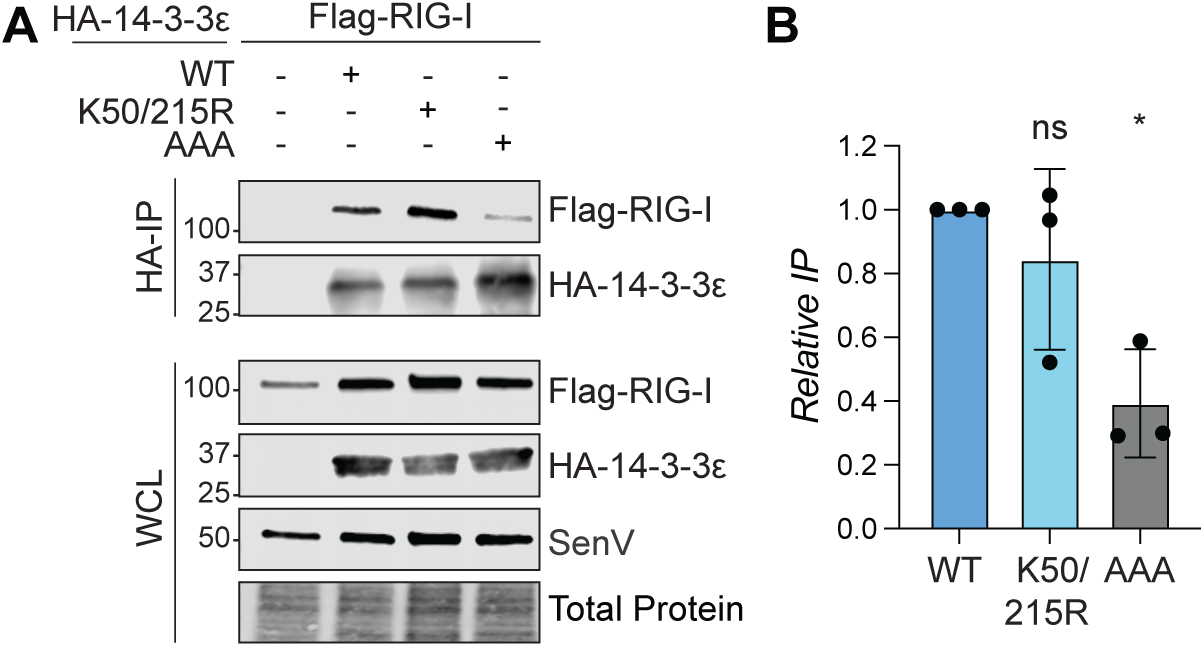
K50 and K215 are not required for 14-3-3ε interaction with RIG-I. (A) Immunoblot analysis of anti-HA immunoprecipitated extracts and inputs from 293T RIG-I KO cells expressing Flag-RIG-I, and HA-14-3-3ε WT, K50R/K215R, or AAA that were SenV-infected (4 h). A representative blot from *n =* 3 independent replicates is shown. (B) Relative quantification of the ratio of RIG-I (IP/relative input) to immunoprecipitated 14-3-3ε (from A) (Mean ± SD, *n =* 3). *p ≤ 0.05 or non-significant (ns) determined by one way ANOVA followed by Tukey’s multiple comparisons test.

### UFMylation of 14-3-3ε differentially regulates MAVS-complex interactions

Since 14-3-3ε facilitates the interaction between RIG-I and MAVS (6), we hypothesized that the UFMylation-deficient 14-3-3ε^K50/215R^ mutant may impair RIG-I-MAVS complex formation. To test this, we performed co-immunoprecipitation assays during SenV infection using a MAVS construct designed to capture CARD-dependent interactions, such as those mediated by the MAVS and RIG-I N-terminal CARDs (63). As expected, overexpression of 14-3-3ε^WT^ enhanced the RIG-I and MAVS interaction compared to the control (Figure 5A-5B). Surprisingly, 14-3-3ε^K50/215R^ did not reduce the association of RIG-I and MAVS, and instead it significantly enhanced this interaction beyond 14-3-3ε^WT^ levels (Figure 5A-5B). This unexpected finding suggests that when 14-3-3ε cannot be UFMylated, RIG-I associates more stably with MAVS but in a potentially non-productive configuration. In contrast to its effect on RIG-I, the K50R/K215R mutation consistently reduced 14-3-3ε interaction with MAVS compared to wild-type (Figure 5A-5B). Together, these results reveal that UFMylation of 14-3-3ε at K50/K215 is associated with MAVS signalosome assembly in an unexpected way: it stabilizes 14-3-3ε-MAVS association while paradoxically dampening RIG-I-MAVS interaction.

**Figure 5.**
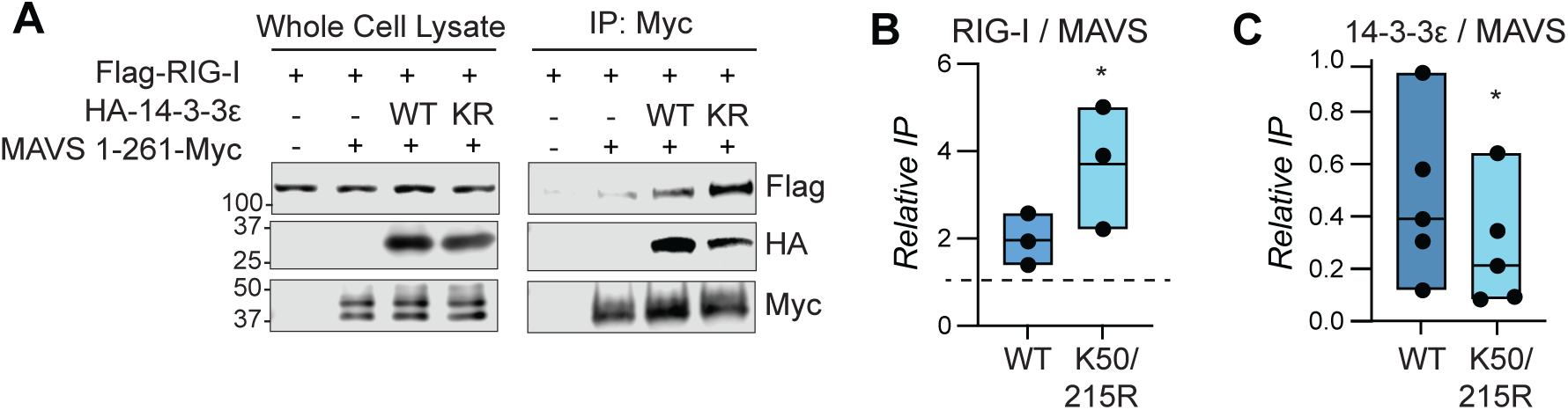
UFMylation of 14-3-3ε differentially regulates MAVS complex interactions. (A) Immunoblot analysis of anti-Myc immunoprecipitated extracts and inputs from 293T RIG-I KO cells expressing MAVS aa1-266-Myc, Flag-RIG-I, and HA-14-3-3ε that were SenV-infected (4 h). A representative blot from *n =* 3 independent replicates is shown. (B) Relative quantification of the ratio of RIG-I (IP/relative input) to immunoprecipitated MAVS. The dashed line indicates the baseline level of RIG-I interaction with MAVS. (C) Relative quantification of the ratio of 14-3-3ε (IP/relative input) to immunoprecipitated MAVS. For (B) and (C), the bars display the data spread, with mean and individual data points shown from independent replicates; *p ≤ 0.05 by ratio paired t test.

## Discussion

Our study identifies UFMylation of 14-3-3ε as a novel regulatory mechanism that promotes RIG-I-mediated innate immune signaling. Upon viral RNA recognition, activated RIG-I must translocate from the cytosol to intracellular membranes to interact with MAVS within a specialized signaling complex termed the “RIG-I translocon.” This complex includes the K63 ubiquitin ligases TRIM25 and Riplet, as well as 14-3-3ε (6, 64). Through biochemical and functional analyses, we demonstrated that UFM1 is covalently conjugated to either lysine residue K50 or K215 of 14-3-3ε as a mono-modification. We further found that mutation of these lysine residues inactivates 14-3-3ε UFMylation and limits the ability of 14-3-3ε to facilitate RIG-I-dependent IFN induction. Surprisingly, we found these UFMylation-inactivating mutations did not affect the interaction between 14-3-3ε and RIG-I. Furthermore, since the 14-3-3ε K50R/K215R mutant retains RIG-I binding comparable to wild-type, the signaling defects we observed cannot be attributed to global disruption of the 14-3-3ε ligand-binding groove. Instead, the K50R/K215R mutations reduced the interaction of 14-3-3ε with MAVS. Our findings reveal that, in addition to facilitating RIG-I translocation to MAVS (6), 14-3-3ε may also associate with MAVS itself, either directly or more likely as part of the RIG-I-MAVS complex. UFMylation of 14-3-3ε stabilizes this 14-3-3ε-MAVS association, as the UFMylation-deficient K50R/K215R mutant shows reduced MAVS complex interaction. Intriguingly, loss of 14-3-3ε UFMylation paradoxically enhances RIG-I-MAVS interaction while simultaneously reducing 14-3-3ε-MAVS binding and impairing downstream signaling. This suggests that UFMylated 14-3-3ε coordinates signaling complex architecture rather than solely promoting RIG-I-MAVS interaction. These data support a model by which UFMylated 14-3-3ε functions beyond simply bringing RIG-I to MAVS; it likely coordinates the proper architecture of the MAVS signalosome ensuring optimal configuration for downstream signal propagation.

Our findings reveal an important distinction between global loss of UFMylation machinery and site-specific inactivation of 14-3-3ε UFMylation. In our previous work, depletion of UFL1 or UFM1 reduced both RIG-I-14-3-3ε and RIG-I-MAVS interactions (21), whereas the current study shows that preventing 14-3-3ε UFMylation through K50R/K215R mutation maintains normal RIG-I-14-3-3ε binding while paradoxically enhancing RIG-I-MAVS interaction. This apparent discrepancy likely reflects the pleiotropic effects of global UFMylation loss versus targeted disruption of a single substrate. The UFMylation machinery likely modifies multiple proteins within the RIG-I signaling pathway, such as other MAVS-associated proteins. Thus, complete loss of the UFMylation machinery would disrupt multiple protein-protein interactions simultaneously, perhaps affecting earlier steps in RIG-I activation and translocation. In contrast, selective loss of 14-3-3ε UFMylation allows other UFMylation-dependent interactions to proceed normally, permitting RIG-I-14-3-3ε complex formation and translocation to MAVS. However, without UFMylated 14-3-3ε to properly coordinate the MAVS signalosome architecture, RIG-I interacts more stably with MAVS. However, our results suggest that this enhanced interaction presumably facilitates a non-productive configuration of the MAVS signalosome as it fails to support robust downstream signaling. This model positions 14-3-3ε UFMylation as a critical quality control checkpoint that ensures productive, rather than merely physical, RIG-I-MAVS complex assembly.

Previously, we demonstrated that UFL1 is recruited to ER-mitochondrial contact sites following RIG-I activation (21, 56). Our current work extends this model by identifying 14-3-3ε as one of the UFL1 target proteins that regulates RIG-I signaling. Interestingly, the UFMylated lysine residues on 14-3-3ε are strategically located within the conserved amphipathic ligand-binding groove, suggesting that UFMylation may introduce negative charges that could alter local electrostatic interactions or induce conformational changes in this region (14). These modifications likely influence how 14-3-3ε coordinates protein partners at MAVS-localized membrane signaling sites. The observation that only mono-UFMylation occurs on 14-3-3ε aligns with structural constraints within the protein: K50 and K215 reside on opposite faces of the ligand-binding groove, which likely makes it structurally feasible for only one site to be UFMylated per molecule (10). While the K50R/K215R mutations abolish UFMylation at these sites, we cannot definitively exclude that these mutations might also disrupt other PTMs. Indeed, K50 in 14-3-3ε and other 14-3-3 isoforms can undergo acetylation, SUMOylation, and ubiquitination (8, 65, 66), suggesting that competing PTMs at the same residue could mediate context-dependent functional switching between cellular pathways. The presence of two potential UFMylation sites may provide redundancy, ensuring that 14-3-3ε can be UFMylated even if one lysine is occupied by another PTM. Nevertheless, the strong correlation between loss of UFMylation and impaired MAVS interaction, combined with our previous findings that UFL1/UFM1 depletion reduces RIG-I-MAVS interaction (21), supports UFMylation as a critical modification that occurs on these residues during RIG-I signaling. Our findings align with emerging evidence that 14-3-3 proteins function as molecular chaperones that prevent client protein aggregation (14). UFMylation may enhance these chaperone activities during RIG-I signaling by regulating MAVS signalosome architecture and optimizing the molecular environment for assembly of a productive MAVS signalosome.

Several important questions remain for future investigation. First, while the K50R/K215R mutations abolish UFMylation, we cannot entirely exclude that these mutations might also prevent other PTMs (65, 66). Definitive proof that UFMylation is sufficient to enhance MAVS engagement would require complementary approaches, such as *in vitro* reconstitution interaction studies with UFMylated 14-3-3ε. Second, the exact molecular function by which UFM1 conjugation on 14-3-3ε promotes functional MAVS signalosome assembly remains to be fully elucidated. Our findings suggest it may modulate interactions with MAVS-associated regulatory proteins or stabilize the architecture of this multi-protein complex at membrane signaling sites (67). Future studies are needed to identify the specific protein partners involved. Third, the molecular events and timing that trigger site-specific UFMylation of 14-3-3ε following RIG-I activation warrant investigation, as does determining whether other 14-3-3 isoforms undergo similar UFMylation-dependent regulation. Finally, while overexpression studies were necessary for dissecting mechanism, validation with endogenous proteins will be important to determine the physiological magnitude, stoichiometry, and occupancy of 14-3-3ε UFMylation following RIG-I activation during viral infection. Such studies await development of improved reagents, as currently available 14-3-3ε antibodies have limited utility for endogenous co-immunoprecipitation studies. Despite these open questions, our findings establish UFMylation of 14-3-3ε as a critical regulatory mechanism in RIG-I signaling and position this modification system as an important node in innate immune regulation.

The regulatory role of UFMylation in RIG-I signaling parallels its function in other immune pathways. Similar to how UFMylation of PD-1 affects T cell activation (42, 44), modification of 14-3-3ε demonstrates how this ubiquitin-like system provides context-dependent tuning of immune responses. By identifying K50 and K215 on 14-3-3ε as critical UFMylation sites that promote RIG-I signaling, our study reveals how this post-translational modification system tunes diverse innate immune pathways. These findings have potential therapeutic implications, as the UFMylation pathway represents a target for modulating antiviral innate immune responses in conditions characterized by either insufficient immunity or excessive inflammation. Moreover, given that multiple viruses have evolved mechanisms to manipulate UFMylation (54, 55), targeting this pathway could potentially disrupt various aspects of viral infection. Understanding the molecular details of UFMylation-dependent immune regulation, as demonstrated here for 14-3-3ε, may ultimately guide development of selective immunomodulatory approaches.

## Data Availability Statement

Plasmids, cell lines, and other reagents described in this study are available from the corresponding author upon reasonable request.

## Acknowledgements

We thank those colleagues who generously provided reagents, as indicated in the Methods, the Duke Functional Genomics Core, the Duke Proteomics and Metabolomics Core Facility, and members of the Horner Lab for valuable feedback and discussion.

## Use of Generative AI

During manuscript preparation, the authors used Perplexity AI (Claude-based model, October 2025 version) to assist with editing, improving clarity, and optimizing sentence structure. All AI-generated suggestions were reviewed, revised, and approved by the authors. The authors take full responsibility for the content of this publication.

## Methods

### Cell lines, viruses, and transfection

293T, 293T-UFM1 KO, 293T-RIG-I KO, and A549-RIG-I KO were grown in Dulbecco’s modification of Eagle’s medium (DMEM; Mediatech) supplemented with 10% fetal bovine serum, 1X minimum essential medium nonessential amino acids, and 25 mM of HEPES buffer. The 293T cells (CRL-3216) were obtained from American Type Culture Collection (ATCC). 293T-UFM1 KO and 293T-RIG-I KO were previously described (21, 59). A549-RIG-I KO were generated by using a purified ribonucleoprotein complex consisting of Cas9 protein and single-guide RNAs (sgRNAs) targeting RIG-I (Synthego). Cas9 protein and sgRNAs were mixed at a ratio of 1:6 and then added to 1x10^6^ A549 cells in Neon Resuspension Buffer R, followed by electroporation using the Neon Transfection System (Invitrogen). Following recovery, single cell clones were isolated and validated by immunoblot. All cell lines were verified as mycoplasma-free using the Look-Out Mycoplasma PCR detection kit (Sigma-Aldrich). Sendai virus Cantell strain was obtained from Charles River Laboratories and used at 200 HAU/ml. Sendai virus infections were performed in serum-free media (30 to 60 min), after which complete medium was added. DNA transfections were performed using FuGene 6 (Promega). Cells were incubated for 24-48 hours post transfection prior to experimental treatment.

### Plasmids

The following plasmids have been previously described: pIFN-β-luc (68), pCMV-Renilla (Promega), pcTak_Myc-5_UFL1 (21), pETDuet-His-TEV-UFL1-StrepII-3C-UFBP1^(21-end)^(gift of Dr. Yogesh Kulathu) (29), pTRIPZ IFNB1^2x^-Gluc (gift of Dr. Nandan Gokhale) (59), and pET-NT*-HRV3CP (gift of Dr. Gottfried Otting; Addgene #162795) (62). The following plasmids were generated by insertion of PCR-amplified fragments into BamHI-to-NotI digested pGEX-GST using InFusion cloning (Takara): pGEX-GST-14-3-3ε^WT^, pGEX-GST-14-3-3ε^K50/215R^, pGEX-GST-UBA5, pGEX-GST-UFC1, and pGEX-GST-UFM1. The following plasmids were generated by site-directed mutagenesis: pEF-TAK-HA-14-3-3ε^K50/215R^, pEF-TAK-HA-14-3-3ε^AAA^ (E251-253A), and pEF-TAK-Flag-UFM1^ΔC2^. The following were cloned by InFusion: pEF-MAVS-1-261-Myc; pEF-TAK-Flag-UFM1^WT^, pEF-TAK-Flag-UFM1^ΔC3^, and pEF-TAK-Flag-UFM1 G83A from pEF-TAK-HA-UFM1 (21); pEF-TAK-Flag-RIG-I from pEF-Bos-Flag-RIG-I (69), pEF-TAK-HA-14-3-3ε^WT^ from pEF-TAK-Flag-14–3-3ε (21), and pEF-TAK-Myc-UFBP1 by PCR from BC115617. Plasmid sequences were verified by DNA sequencing.

### IFN-β promoter luciferase assay

IFN-β promoter luciferase assays were performed at 18 to 24 hours after treatment and transfection of pIFN-β-luc and pCMV-Renilla by using the Dual-Luciferase Reporter System (Promega). *Firefly* luciferase was normalized to *Renilla* luciferase, whose expression is driven by the CMV promoter. For the *Gaussia* luciferase assays, 48 hours post transfection of pTRIPZ IFNB1^2x^-Gluc, supernatant was assayed for *Gaussia* luciferase using the *Renilla* Luciferase Assay System (Promega). All samples were read using a BioTek Synergy2 microplate reader.

### Immunoblotting

Cells were lysed with either RIPA (10 mM Tris HCl pH [7.5], 150 mM NaCl, 0.5% sodium deoxycholate, and 1% Triton X-100) or NP-40 Lysis Buffer (20 mM Tris HCl [pH 7.4], 100 mM NaCl, 0.5% NP-40) supplemented with protease inhibitor mixture (Sigma-Aldrich), and Halt inhibitor phosphatase (Thermo Fisher Scientific), and lysates were isolated by centrifugation. Quantified protein was resolved by SDS/PAGE and transferred to nitrocellulose membranes. Membranes were stained with Revert 700 total protein stain (Li-COR Biosciences) and then blocked with 3% bovine serum albumin in phosphate-buffered saline containing 0.01% Tween-20 (PBS-T). Following incubation with primary antibodies, membranes were incubated with species-specific horseradish-peroxidase (HRP)-conjugated antibodies (Jackson Immuno Research 1:5000), followed by membrane treatment with Clarity Western ECL substrate (Bio-Rad) and imaging on a LICOR Odyssey FC. The following antibodies were used for immunoblotting: R-anti-SenV (MBL, 1:1000), HA-HRP (Sigma-Aldrich 1:1000-1:3000), Flag-HRP (1:1000-1:5000), and M-anti-Myc (Cell Signaling Technology 1:1000).

### Protein immunoprecipitation

Cells were lysed with either RIPA (10 mM Tris HCl [pH 7.5], 150 mM NaCl, 0.5% sodium deoxycholate, and 1% Triton X-100) or NP-40 Lysis Buffer (20 mM Tris HCl [pH 7.4], 100 mM NaCl, 0.5% NP-40) supplemented with protease inhibitor mixture (Sigma-Aldrich), and Halt phosphatase inhibitor, and 10% glycerol. Quantified protein (between 100-200 µg) was incubated anti-HA magnetic beads (Invitrogen) in lysis buffer either at room temperature for 1-2 hours or 4°C overnight with head-over-tail rotation. Beads were washed 3X in lysis buffer and eluted with 2X Laemmli buffer (Bio-Rad) with 5% 2-mercaptoethanol at 95°C for 5 min. Proteins were resolved by SDS/PAGE and immunoblotting, as above. For UFMylation precipitations, cells were lysed UFMylation lysis buffer (50 mM Tris [pH 8], 150 mM NaCl, 0.5% sodium deoxycholate, and 1% NP-40) supplemented with 10% glycerol, 1:100 protease and phosphatase inhibitors, as above, and 1:100 *N*-ethylmaleimide (10 mM final) (Sigma-Aldrich). Post-nuclear lysates were boiled at 95°C for 10 min and incubated with Pierce anti-HA magnetic beads with UFMylation lysis buffer. Beads were washed 3X in high salt buffer (10 mM Tris [pH 8], 1 mM EDTA, 500 mM NaCl) and eluted with 2X Laemmli buffer with 5% 2-mercaptoethanol at 95°C for 5 min.

### Quantification of immunoblots

Immunoblots were imaged using the LICOR Odyssey FC system and quantified using ImageStudio software, and raw values were normalized to relevant controls for each antibody. RIG-I and 14-3-3ε interaction was quantified as ratio of the RIG-I levels in the IP relative to the input normalized to the 14-3-3ε pull down. RIG-I or 14-3-3ε interaction with MAVS was quantified as ratio the RIG-I or 14-3-3ε levels relative to the input normalized to the MAVS pull down. The RIG-I interaction with MAVS was normalized to the vector control (shown as a dashed line) to display the relative enhanced interaction in the presence of 14-3-3ε, while the 14-3-3ε interaction with MAVS shows raw values normalized to display the interaction differences between conditions.

### Purification of GST/His-tagged proteins

pET28A-His-UBA5, pET28A-His_6_-UFC1, pET28A-His_6_-UFM1^dSC^, pGEX-GST-14-3-3ε^WT^, pGEX-GST-14-3-3ε^K50/215R^, and pETDuet-His_6_-TEV-UFL1-StrepII-3C-UFBP1^(21-end)^ were transformed in *E. coli* Rosetta cells. Cells were cultured at 37°C until OD600 reached 0.6 to 0.8. Then, cells were induced with a final concentration of 1 mM IPTG. Cells were further cultured at 30°C for 3.5-4 hours. Cells were collected by centrifugation for 30 min at 7,000 rpm. The pellets were lysed in NET-N buffer (50 mM Tris-HCl [pH 7.5], 150 mM NaCl, 0.5% IGEPAL, supplemented with Aprotinin, Leupeptin, PMSF, and dithiothreitol (DTT) at 1:100. Cells were sonicated 3 times for 30 sec at output 6-7 and a duty cycle of 80%. Then, lysates were centrifuged for 30 min at 12,000 rpm at 4°C. Supernatant was incubated with GST beads that were washed with NET-N buffer with rotation at 4°C for 1-2 hours. The beads were washed 3 times with NET-N buffer and 3 times with PBS supplemented with 1:100 DTT. After washes, GST tag was cleaved with purified human rhinovirus 3C protease (purified as described (62)) and protein was eluted with PBS by incubation with rotation at 4°C overnight.

### *In vitro* UFMylation

2 µM of substrate, 0.25 µM of UBA5, 5 µM UFC1, 20 ng UFL1/UFBP1, 30 µM UFM1 were incubated in a reaction buffer containing 50 mM HEPES [pH 7.5], 10 mM MgCl_2_, with or without 5 mM ATP for 1 hour at 37°C. The reaction was stopped by the addition of 2X SDS loading with 5% 2-mercaptoethanol and incubated at 95°C for 5 min. The reaction products were then separated by SDS/PAGE and analyzed by immunoblotting, as above.

### RNA Analysis

Total cellular RNA was extracted using RNeasy Plus mini kit (Qiagen). RNA was then reverse transcribed using iScript complementary DNA (cDNA) synthesis kit (Bio-Rad), following manufacturer’s instructions. The resulting DNA was diluted 1:3 in double distilled water. RT-qPCR was performed in triplicate using the Power SYBR Green PCR master mix (Thermo Fisher Scientific) and QuantStudio 6 Flex RT-PCR system. Oligonucleotide sequences are as follows: IFNB1 F-5’-CTTTGCTATTTTCAGACAAGATTCA; R-5’GCCAGGAGGTTCTCAACAAT, IFNL1 F-5’-CTTCCAAGCCCACCACAACT; R-5’-GGCCTCCAGGACCTTCAGC, and GAPDH F-5’-AAGGTGAAGGTCGGAGTCAAC; R-5’-GGGGTCATTGATGGCAACAATA.

### Sample Preparation and LC-MS/MS Analysis

Samples were spiked with undigested bovine casein at a total of either 120 or 240 pmol as an internal quality control standard. Next, samples were supplemented with 10 μL of 20% SDS, reduced with 10 mM DTT for 30 min at 80°C, alkylated with 20 mM iodoacetamide for 30 min at room temperature, then supplemented with a final concentration of 1.2% phosphoric acid and 546 μL of S-Trap (Protifi) binding buffer (90% MeOH/100 mM TEAB). Proteins were trapped on the S-Trap micro cartridge, digested using 20 ng/μL sequencing grade trypsin (Promega) for 1 hour at 47°C, and eluted using 50 mM TEAB, followed by 0.2% FA, and lastly using 50% ACN/0.2% FA. All samples were then lyophilized to dryness. Samples were resolubilized using 48 μL of 1% TFA/2% ACN with 25 fmol/μL yeast ADH. Quantitative LC/MS/MS was performed on 1.5 μL (3.13% of total sample) using an MClass UPLC system (Waters Corp) coupled to an Orbitrap Fusion Lumos high resolution accurate mass tandem mass spectrometer (Thermo Fisher Scientific) via a nanoelectrospray ionization source. Briefly, the sample was first trapped on a Symmetry C18 20 mm × 180 μm trapping column (5 μl/min at 99.9/0.1 v/v water/acetonitrile), after which, the analytical separation was performed using a 1.8 μm Acquity HSS T3 C18 75 μm × 250 mm column (Waters Corp.) with a 90-min linear gradient of 5 to 30% acetonitrile with 0.1% formic acid at a flow rate of 400 nanoliters/min with a column temperature of 55°C. Data collection on the Fusion Lumos mass spectrometer was performed in a data-dependent acquisition (DDA) mode of acquisition with a r=120,000 (@ m/z 200) full MS scan from m/z 375 – 1500 with a target AGC value of 4e5 ions was performed. MS/MS scans were acquired in the linear ion trap in “rapid” mode with a target AGC value of 1e5 and max fill time of 100 ms. The total cycle time 2s, with total cycle times of 2 sec between like full MS scans. A 20s dynamic exclusion was employed to increase depth of coverage. The total analysis cycle time for each sample injection was approximately 2 hours.

### Proteomics Data Analysis

Following UPLC-MS/MS analyses, data were imported into Proteome Discoverer 2.5 (Thermo Fisher Scientific). In addition to quantitative signal extraction, the MS/MS data was searched against the SwissProt *H. sapiens* database (downloaded in Nov 2019) and a common contaminant/spiked protein database (bovine albumin, bovine casein, yeast ADH, etc.), and an equal number of reversed-sequence “decoys” for false discovery rate determination. Sequest (v 2.5, Thermo PD) was utilized to produce fragment ion spectra and to perform the database searches. Database search parameters included fixed modification on Cys (carbamidomethyl) and variable modification on Met (oxidation) and Lys (UFMylation). Search tolerances were 2ppm precursor and 0.8Da product ion with semi-trypsin enzyme rules. Discoverer were used to annotate the data at a maximum 1% protein false discovery rate based on q-value calculations. Note that peptide homology was addressed using razor rules in which a peptide matched to multiple different proteins was exclusively assigned to the protein has more identified peptides. Protein homology was addressed by grouping proteins that had the same set of peptides to account for their identification. A master protein within a group was assigned based on % coverage.

### Statistical analysis

We used ratio paired t test or one-way ANOVA followed by appropriate multiple comparisons test using GraphPad Prism software. Graphed values are presented as mean ± SD (n = 3) or as individual data points with spread; \**P* ≤ 0.05, \*\**P*≤0.01, and \*\*\**P* ≤0.001.

